# Large-scale miRNA-Target Data Analysis to Discover miRNA Co-regulation Network of Abiotic Stress Tolerance in Soybeans

**DOI:** 10.1101/2021.09.09.459645

**Authors:** Haowu Chang, Tianyue Zhang, Hao Zhang, Lingtao Su, Qing-Ming Qin, Guihua Li, Xueqing Li, Li Wang, Tianheng Zhao, Enshuang Zhao, Hengyi Zhao, Yuanning Liu, Gary Stacey, Dong Xu

**Author notes:** Corresponding author: Hao Zhang, Dong Xu. These authors contributed equally to this work (Haowu Chang, Tianyue Zhang). H.Z. and D.X. conceived and designed the study; L.S., Q.Q., G.L., X.L., T.H.Z. and L.W. assembled the data; H.C. and T.H.Z. performed the analyses; H.C. wrote the modeling code; Y.L., H.W. and G.S. assisted with interpretation of the results; H.C. and T.Y.Z. wrote the manuscript; H.Z., D.X. and G.S. reviewed and revised the manuscript. The authors declare no competing interests.

## Abstract

Although growing evidence shows that microRNA (miRNA) regulates plant growth and development, miRNA regulatory networks in plants are not well understood. Current experimental studies cannot characterize miRNA regulatory networks on a large scale. This information gap provides a good opportunity to employ computational methods for global analysis and to generate useful models and hypotheses. To address this opportunity, we collected miRNA-target interactions (MTIs) and used MTIs from Arabidopsis thaliana and Medicago truncatula to predict homologous MTIs in soybeans, resulting in 80,235 soybean MTIs in total. A multi-level iterative bi-clustering method was developed to identify 483 soybean miRNA-target regulatory modules (MTRMs). Furthermore, we collected soybean miRNA expression data and corresponding gene expression data in response to abiotic stresses. By clustering these data, 37 MTRMs related to abiotic stresses were identified including stress-specific MTRMs and shared MTRMs. These MTRMs have gene ontology (GO) enrichment in resistance response, iron transport, positive growth regulation, etc. Our study predicts soybean miRNA-target regulatory modules with high confidence under different stresses, constructs miRNA-GO regulatory networks for MTRMs under different stresses and provides miRNA targeting hypotheses for experimental study. The method can be applied to other biological processes and other plants to elucidate miRNA co-regulation mechanisms.

## Introduction

The growth and development of crops are often restricted due to various environmental stresses, which can lead to poor harvests and yields below their genetic potential [1, 2]. In the past decade, microRNAs (miRNAs) have been identified as important gene expression regulatory factors that play an essential role in plant growth and development [3]. miRNA can target multiple genes, and multiple miRNAs can also target the same gene. miRNAs are involved in the expression of stress-responsive genes and the plant’s ability to adapt to environmental change [4]. Different stresses can induce differential expressions of corresponding miRNAs in plants, while some miRNAs can respond to several abiotic stresses simultaneously [5-7]. Therefore, studying the cooperative relationship among miRNAs and the interactions with their target genes is important for understanding the role of miRNAs in controlling plant growth and development.

miRNAs may respond to adverse effects on plant growth and development, such as drought, salinity, temperature and other abiotic environmental factors. It was shown that willow leaves exposed to drought or high temperature induce differential expressions of some miRNAs [8]. For example, miR169c plays a negative regulatory role under drought stress by inhibiting the expression of its target gene nuclear factor Y-A (NF-YA) [9]. miR172a [10] and miR172c [11] endow plants with a tolerance to salt stress and water deficiency. Meanwhile, miRNAs also indirectly respond to abiotic stress by regulating other biological macromolecules. For example, miR398c can negatively regulate multiple peroxisome-related genes (GmCSD1a/b, GmCSD2a/b/c and GmCCS) and affect the drought tolerance of the soybean [12]. miR399a/b participates in developing resistance to salt stress [13]. miR5036, miR862a and miR398a/b respond to phosphorus starvation and salt stress [14]. Furthermore, miR166k/o, miR390g, and miR396c/k mediate BX10 (Al-tolerant genotype) root elongation, and miR169r triggers the BD2 (Al-sensitive B genotype) oxidative stress, which in turn triggers a different type of plant aluminum tolerance between BX10 and BD2 [15]. This indicates that miRNA may regulate plant growth under abiotic stress through a complex network. However, current studies typically explore the role of a single miRNA in response to abiotic stresses. How multiple miRNAs work together as a co-regulatory mechanism has not been significantly explored.

Several studies have uncovered interesting miRNA interactions. For example, it was shown that miRNA lin-4 and let-7 act cooperatively in Drosophila [16], while miR375, miR124 and let7b coordinately regulate Mtpn expression in mammals [17]. Similarly, the miRNAs, miR125a, miR125b and miR20, work together to down-regulate erbB2/erbB3 in breast cancer cells [18,19]. Furthermore, compared with individual miRNAs alone, the combination of miR16, miR34a and miR106b results in a stronger immortalized human mammary epithelial cell cycle arrest [20]. It was also found that miR21, miR23a and miR27a are three cooperative repressors in a network of tumor suppressors [21]. The same combination effects also exist in plants. For example, miR160 and miR167 are involved in the adventitious root program of Arabidopsis [23]. miR156 and miR172 play a role in the transition of soybean nutrition [24], and miR172 also plays a role in flowering by inhibiting AP2 [25]. Transgenic studies of miR482, miR1512 and miR1515 showed that their over-expression may lead to a substantial increase in the number of soybean nodules [26]. Another study verified networks of 365 tissue-specific miRNA-target interactions (MTIs) [27]. The synergistic effects of miRNAs provide a new systematic perspective for the entire microRNome [23], which calls for a global analysis of miRNA-target interactions.

With the growing miRNA-target data, several methods have been developed and applied to explore this field. Shalgi et al. first constructed a miRNA network from the target genes predicted by PicTar and TargetScan [28]. Xu et al. constructed a human miRNA-miRNA functional synergy network through co-regulation functional modules [29]. Meanwhile, biclustering was also applied for two different types of objects (gene and miRNA in this case) belonging to the same cluster. In the past two decades, various bi-clustering methods have been developed [30,44-46]. SAMBA [31], ISA [32], BIMAX [33], QUBIC [34] and FABIA [35] are some commonly used general algorithms. Contiguous column coherent (CCC) biclustering [36-41] and LateBiccluster [39] are designed for temporal data analysis. BicPAM [40, 41], BicNET [42] and MCbiclust [43] are the latest tools. Gianvito et al. applied the biclustering algorithm to predict human miRNA-mRNA modules [47]. The application of biclustering algorithms and miRNA-target regulation module (MTRM) mining is feasible and important for analyzing miRNA regulation mechanisms. Compared with traditional clustering methods, such as Bimax [103] and BiBit [69], CUBiBit [57] shortened the computing time and provided an optimized method for finding modules in larger data. However, the result obtained by CUBiBit was mostly a fully-connected bipartite graph, and the relationship between miRNA and the target gene is complex and interactive.

In this study, we analyzed the relationship between the miRNA regulatory modules in response to abiotic stresses in the soybean as a means for extending our understanding of soybean resistance mechanisms. Previously, Xu et al. provided a soybean miRNA-gene network, SoyFN, based on predicted miRNA targets [49]. However, this work was based only on sequence comparisons, which may result in a high false discovery rate. In contrast, in our work, we collected experimentally proven miRNA-target relationships based on degradome sequencing in the soybean, as well as the stringent homologs of miRNA-target pairs in *Arabidopsis thaliana* and *Medicago truncatula*. Based on these reliable miRNA-target data, we performed a biclustering analysis, and iteratively fused the overlapping biclusters based on the SoyNet network to obtain the soybean miRNA-target regulatory modules in response to abiotic stresses. We provide soybean MTRMs with high confidence relevant to various stresses and present the miRNA-GO regulatory networks of these modules. Capturing these miRNA-target modules with biological significance expands our understanding of the complex regulatory mechanisms of miRNA. The methods used should be easily applicable to other plant and animal systems where sufficient data exists to perform the analyses.

## Results

We obtained 90,064 confirmed soybean MTIs based on multiple experimental data sources and 1,189 potential soybean MTIs based on homology to experimental data from *Arabidopsis thaliana* and *Medicago truncatula*. A multi-level iterative bi-clustering analysis resulted in 483 soybean miRNA-target regulatory modules. We analyzed the enrichment results of the top five biclusters. In addition, we identified 37 abiotic stress-related modules and predicted the underlying miRNA regulatory pathway networks.

### Identification of MTIs

We collected soybean miRNA-target data based on databases and related publications. First, we gathered all the soybean miRNA-target interactions (MTIs) verified by degradome sequencing and biological experiments by mining published data, including 14,958 pairs of MTIs from DPMIND [50], 86,427 pairs of MTIs from starBase [51], and 38 pairs of MTIs from TarBase [52]. Additional data came from fourteen publications [70-83], as shown in Supplemental Table 1 (Supplemental Table S1). A total of 111,650 pairs of soybean MTIs were obtained. After removing 21,586 redundant pairs of MTIs, 90,064 pairs remained.

To expand MTIs, we predicted the target relationship between potential miRNAs and targeting genes from the MTIs of *Arabidopsis thaliana* and *Medicago truncatula* based on homology. For Arabidopsis MTIs, 43 pairs were collected from TarBase, 66,212 pairs from starBase, 4682 pairs from DPMIND, 106 pairs from miRTarBase, and 369 pairs from 5 related publications [84-88], resulting in a total of 71,412 pairs of Arabidopsis MTIs, in which 12,094 MTIs became unique after removing 59,318 pairs of redundant MTIs. For Medicago MTIs, there were 302 pairs from TarBase, 22,010 pairs from starBase, 781 pairs from DPMIND, and 1,349 pairs from three publications [89-91], resulting in a total of 24,442 pairs of MTIs, in which 4,394 MTIs became unique after removing 20,048 pairs of redundant MTIs. There are 33 miRNAs with identical sequences between Arabidopsis and the soybean, covering 3,415 targeting genes in Arabidopsis and 4,190 genes in the soybean. There are also 49 miRNAs with identical sequences between Medicago and the soybean, covering 1,180 targeting genes in Medicago and 5,096 genes in the soybean. We removed any redundant MTIs resulting from these. We further validated homology-based MTIs using three miRNA target prediction tools that performed well in general plants, i.e., psRNAtarget [53, 54], TAPIR [55], and Targetfinder [56]. The relationship between a specific miRNA and its target was validated by at least one of the prediction tools. In the Arabidopsis derived MTIs, 943 pairs of MTIs were validated in the psRNAtarget, 152 pairs in TAPIR, and 158 pairs in Targetfinder, confirming a total of 961 unique pairs of MTIs. In the Medicago-derived MTIs, 968 pairs of MTIs were validated in the psRNAtarget, 144 in TAPIR, and 134 in Targetfinder, confirming a total of 986 unique pairs of MTIs, as shown in Supplemental Figure S1. There is a high overlap between the two sets of MTIs (Supplemental Table S2). After removing the redundant ones, a total of 1,189 pairs were used to expand soybean MTIs.

### miRNA-target regulatory modules

We integrated the 90,064 soybean MTIs with the 1,189 MTIs based on homology. We removed MTIs involving genes that do not have the glyma2 ID. A total of 11,018 MTIs were removed, and the remaining 80,235 MTIs were used for analysis in the following tasks.

We applied CUBiBit [57] for bi-clustering analysis, with the smallest scale 2×2 or 6×2 for miRNA-target modules (i.e., at least two or six target genes and at least two miRNAs in each module), resulting in 15,380 (2×2) miRNA-target modules or 2,461 (6×2) miRNA-target modules. We contracted the overlapping modules using a multi-level iterative fusion method based on the soybean gene relationship network (See Methods), yielding 6,577 (2×2) and 812 (6×2) soybean miRNA-target regulatory modules after removing the modules that were completely included in the preliminary clustering module.

We next merged MTRMs according to the set threshold until the level converged stably (level represents the number of iterations). The iterative fusion of each level is shown in Figure 1. We compared the iterative results at different scales. Soybean MTRMs at the 2×2 scale showed better results at level 10, which contains 2,715 MTRMs. Soybean MTRMs at the 6×2 scale showed a better effect at level 7, which contains 483 MTRMs. Comparing the cluster score based on the GO calculation between the two scales of stable convergence (Figure 1c) shows that the cluster score quality at the 6×2 scale is higher than that at the 2×2 level (Supplemental Table S3). Hence, we used the GO enrichment analysis result on 483 soybean MTRMs obtained at the 6×2 level 7.

**Figure 1.**
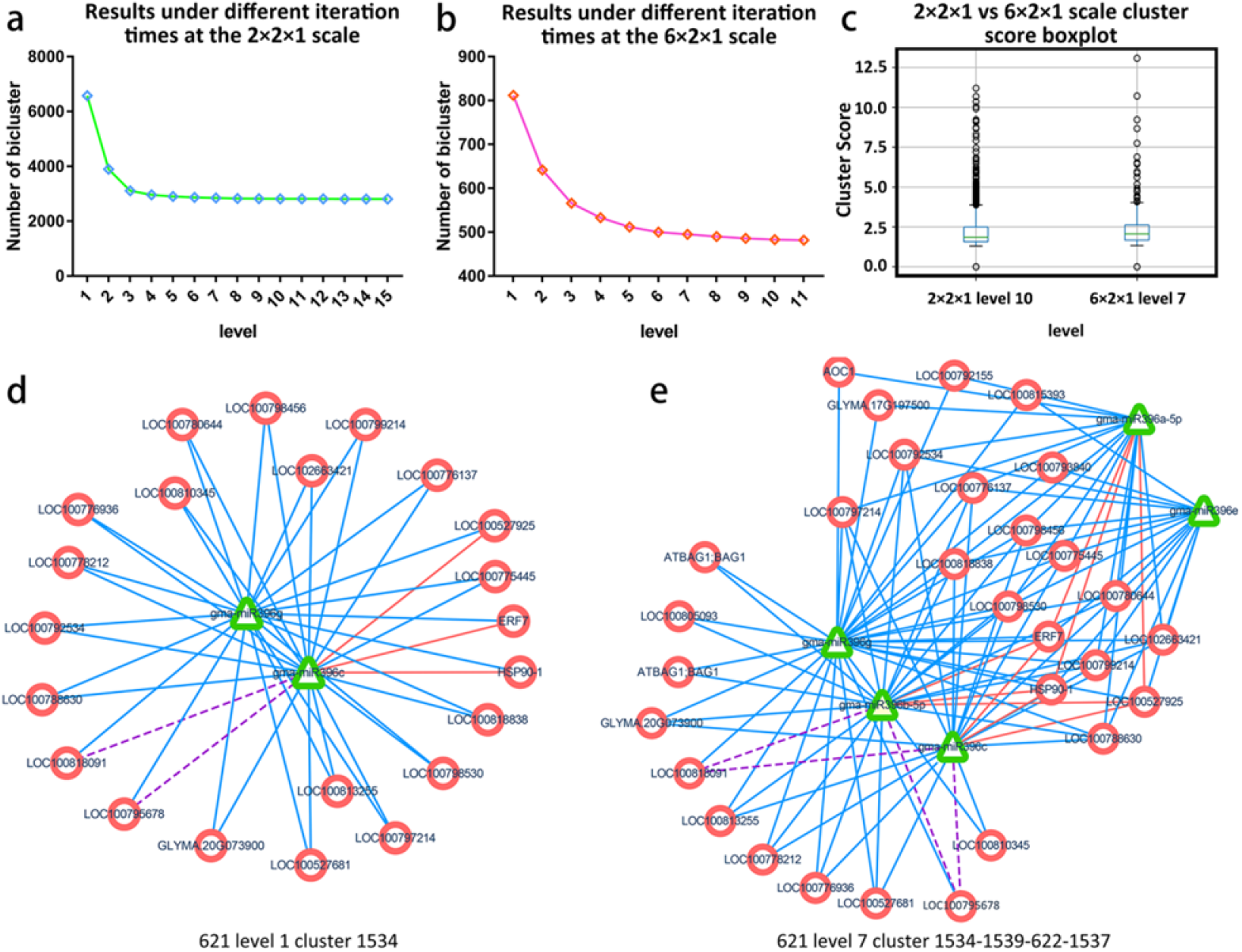
Multi-level iterative biclustering results of soybean MTRMs. (a) Results under different iteration times at the 2×2 scale, (b) results under different iteration times at the 6×2 scale, and (c) the boxplot of the cluster score is calculated based on the gene ontology (GO) under the two scales when converging to a stable level, where based on the overall distribution, the results at the 6×2 scale are better; (d) shows the MTRM bicluster at level 1 before the 6×2 scale fusion, and (e) shows the corresponding MTRM bicluster at level 7 after the 6×2 scale fusion.

To compare the MTRMs before and after fusion, we extracted an MTRM bicluster, as shown in Figure 1(e) from the level 7 clustering results of the 6×2 scale and plotted it with the corresponding MTRMs under level 1 before the fusion, as shown in Figure 1(e) and after Figure 1(d), which is a level-7 fusion. The module (1534) is at level 1 before the fusion has 2 miRNAs and 22 targeted genes. At level 7, the module (1534) fused an additional three modules 1539, 622, and 1537, and each contains miR396. From the perspective of targeting, the module at level 7 has more miRNA-target interactions than the one at level 1.

### Gene ontology (GO) analysis of MTRMs

We screened 254 GO pathways whose GO biological processes (BP) satisfied the p-value < 0.00001 for the GO enrichment and obtained 483 soybean MTRMs at the 6×2 scale at level 7. We analyzed the relationship among the enriched GO terms through REVIGO [122] with a parameter of 0.5. These GO pathways have a specific aggregation (Figure 2a). MTRMs obtained from a global perspective have several concentrated distributions of GO functions, such as cellular processes, primary metabolism, and cell adhesion, as well as hormone response and negative regulation of biological processes. In addition, there are metachronous positive growth regulations and chalcone biosynthesis. Chalcone plays an important role in soybeans and is involved in the multi-branch pathway of flavonoids and isoflavone biosynthesis [92]. The enrichment results mainly involve positive regulation of development, heterochronic, chalcone biosynthesis, defense response, mitochondrial mRNA modification, sulfate transport, plant-type primary cell wall biogenesis, and cofactor biosynthesis, as shown in Figure 2(b).

**Figure 2.**
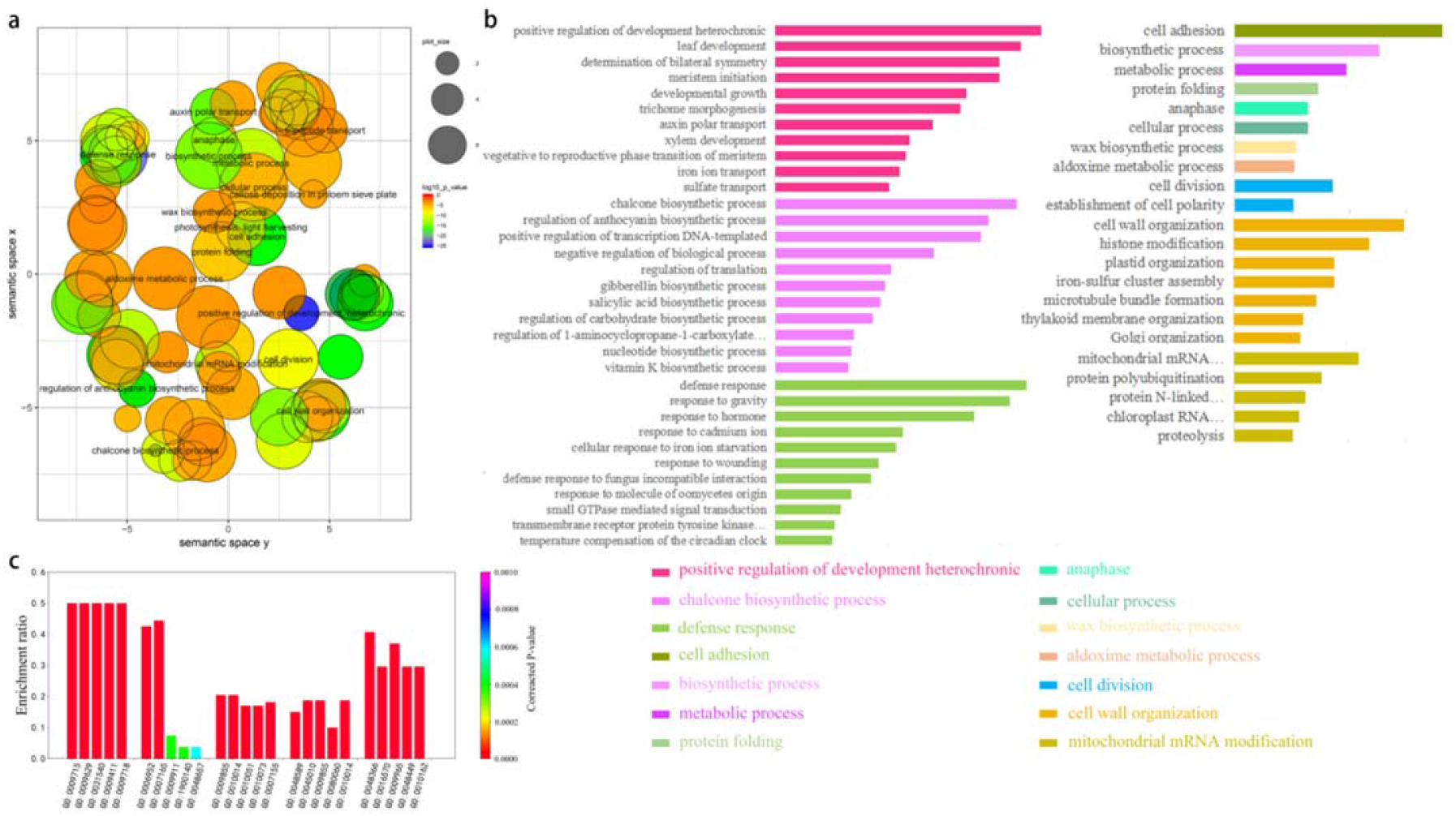
GO analysis of soybean MTRMs. (a) Semantic relevance of GO terms wherein the GO pathway has a certain concentration. (b) GO annotation enriched with 483 soybean MTRMs with enrichment results, which mainly involve positive regulation of development, heterochronic, chalcone biosynthesis, defense responses, and mitochondrial mRNA modification, and (c) GO enrichment of the top five soybean MTRMs. The listed GO terms were enriched with significant p-values < 0.00001

In addition, we extracted the enrichment results of the top biclusters in terms of cluster score among the 483 MTRMs and selected the top five GO terms of each module, as shown in Figure 2(c) and Supplemental Table S4).

### Abiotic stress-related modules

To explore the biological significance of soybean MTRMs, we collected related soybean miRNAs, which revealed 10 types of abiotic stresses, involving 1) drought, 2) salt, 3) cold, 4) nodulation, 5) Pi, 6) oil, 7) rust, 8) soybean isoflavone, 9) pollen development, and 10) phosphorus deficiency based on publications [22, 48, 105-121]. The function annotations of these miRNAs are shown in Supplemental Dataset S1. In most processes, soybean miRNA responses are involved in multiple abiotic responses, as shown in Figure 3(a). Therefore, mining the potential cooperative regulatory modules of these miRNAs is important to understand their role in modulating soybean stress responses.

**Figure 3.**
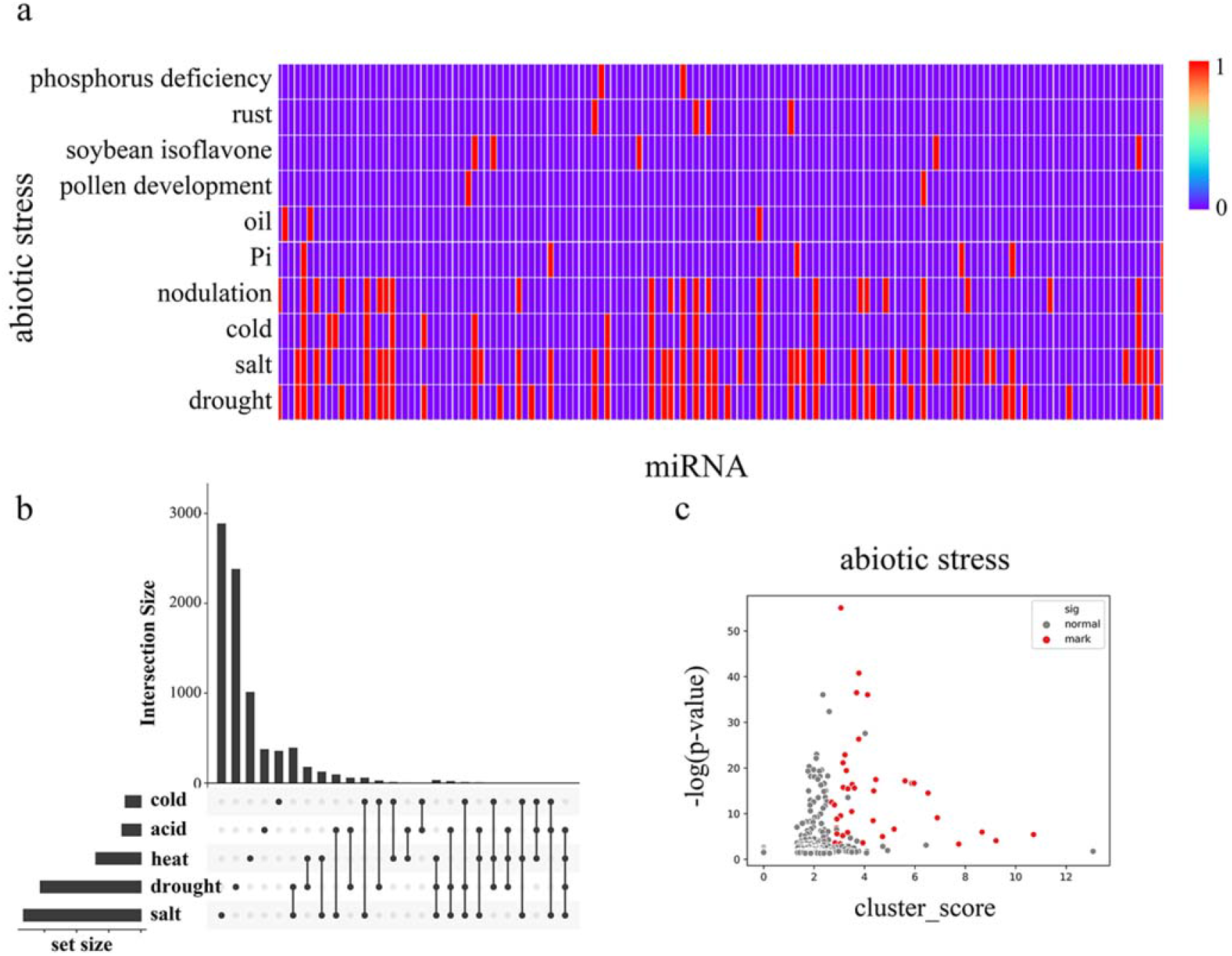
Collected miRNA data on soybeans involved in various abiotic stress responses based on the data statistics from the literature. (a) The distribution of stress types in miRNAs where each vertical line represents one miRNA, and red is marked as relevant, (b) UpSet diagram [123] of modular genes under various abiotic stresses within the horizontal correspondence, where dots are used to refer to the corresponding cold stress, acid stress, heat stress, drought stress, and salt stress on the left. The point-to-point connection is realized longitudinally to indicate the intersection between the corresponding data sets, and the upper bar graph indicates the number of genes in the intersection. In (c), the differentially expressed genes in each MTRM under abiotic stress are shown after screening. We used three indicators to filter the candidate clusters. According to the p-value, the related miRNA purity, and the cluster score of each MTRM gene are placed under the corresponding stress. We selected the corresponding threshold and obtained the stress-related MTRMs with higher reliability and marked them as red dots in Figure 3(c). Supplemental Figure S2 shows MTRMs under other types of stress.

We correlated the 483 soybean MTRMs obtained by clustering with the functional annotations. We selected miRNAs that responded to drought resistance, salt resistance, heat stress, cold stress, acid stress, and performed a statistical analysis on the miRNAs in each of the 483 biclusters. We collected data on the differential expression of soybean genes in the MTRMs under drought, salt, low temperature, cold, and acid stress. Among them, the drought stress data are from GSE76636, the cold stress data are from GSE117686, the acid stress data are from GSE75575, and the salt stress expression data are from the data under 0-h and 12-h treatments of salt stress in GSE57252 (Supplemental Dataset S2). The conditions for screening differentially expressed genes are log2FC > 1, p < 0.05. We obtained 2,145 differentially expressed genes under soybean drought and 1,752 differentially expressed genes under salt treatment. Figure 3(b) shows the genes in the module together with an abiotic stress diagram. At the same time, we calculated the p-values and FDR. We used the Benjamin Graham formula to correct the p-value of the genes in each MTRM for the differentially expressed genes under abiotic stress scenarios through the hypergeometric distribution, as shown in Figure 3(c).

Subsequently, we screened MTRMs related to abiotic stress, drought, and salt stress according to the p-value of differentially expressed genes corresponding to the stress in the MTRMs (p < 0.001, single adversity 0.01), the proportion of the corresponding miRNA family function (miR function ratio), and the cluster score (cluster score > median). The screening results are shown in Supplemental Dataset S2. We obtained 37 MTRMs related to abiotic stress, including 34 MTRMs related to drought stress, 27 MTRMs related to salt stress, 3 MTRMs related to cold stress, and 21 MTRMs related to heat stress. Figure 4(a) shows the set relationship of MTRMs involved in a variety of stresses. The data suggest that soybean miRNAs have basic and universal functional modules in their response mechanisms to drought, high salt, high temperature, low temperature, and other abiotic stresses. There are two shared modules (31 and 493), involving 6 miRNAs and 11 miRNAs, respectively. The 6 miRNAs of module 31 belong to the miR156 family. The regulated gene-enriched GO pathway is a transcription regulation, DNA-dependent (p-value 4.24 e-10), and a vegetative phase change regulation with a p-value of 9.24 e-07. The 11 miRNAs of module 493 are mainly in the miR172 family, in addition to miR156, miR1533, miR4374, miR5782, miR3939. The regulated gene-enriched GO pathway involves an oxidation-reduction process (p-value = 4.66 e-12) and a root hair elongation (p-value = 1.63e-08). Moreover, we also found stress-specific regulatory modules in our results, including 14 drought-specific MTRMs, seven salt-specific MTRMs, and two heat-specific MTRMs (Supplemental Dataset S3).

**Figure 4.**
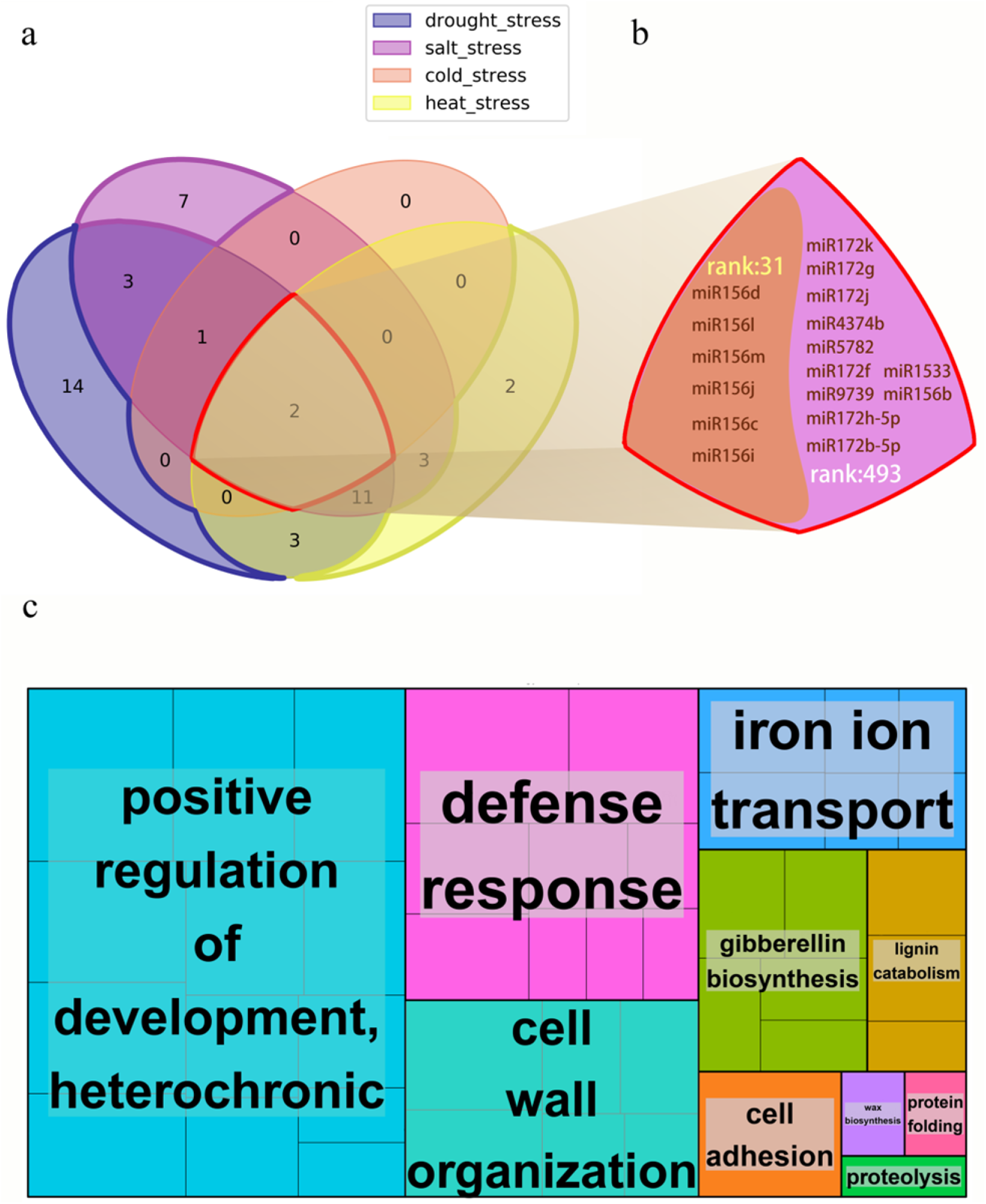
Soybean MTRMs under various abiotic stresses (a) Venn diagram of 37 kinds of soybean MTRMs under various abiotic stresses, including 14 drought-specific MTRMs, seven salt-specific MTRMs, two heat-specific MTRMs, and two shared MTRMs, (b) the miRNAs in the two shared MTRMs 31 and 493, and (c) a GO Treemap of 37 MTRMs under abiotic stress.

The functions of related miRNA regulatory modules under abiotic stress are mainly concentrated in positive regulation of developmental heterochrony, defense responses, cell wall organization, and other biological processes, as shown in Figure (4b). The data of the top-5 modules are shown in Supplemental Table S5.

### miRNA regulatory pathway network under abiotic stress

We explored the regulatory pathway network corresponding to the miRNA of the miRNA-target regulatory module in soybean abiotic stress and analyzed the GO terms of MTRM genes under various abiotic stresses. Stringent screening conditions were used, i.e., the p-value of MTRM stress is 0.001, and the GO BP pathway was selected with a p-value of less than 10^−5^. The REVIGO-based GO language correlation analysis is shown in Figure 5. GO channels with similar functions are closer in distance in the figure. The small RNAs targeting a certain cluster of GO are functionally close, and the 37 MTRMs of soybeans in abiotic stresses identified in this study mainly focus on resistance response, iron transport, positive growth regulation, and cell wall organization. Under abiotic stress, the cooperating miRNA regulatory modules of the soybean mainly regulate these pathways to respond to the stress environment. The correlation analysis between drought stress and salt stress with specificity is shown in Figure 5 (b) and (c).

**Figure 5.**
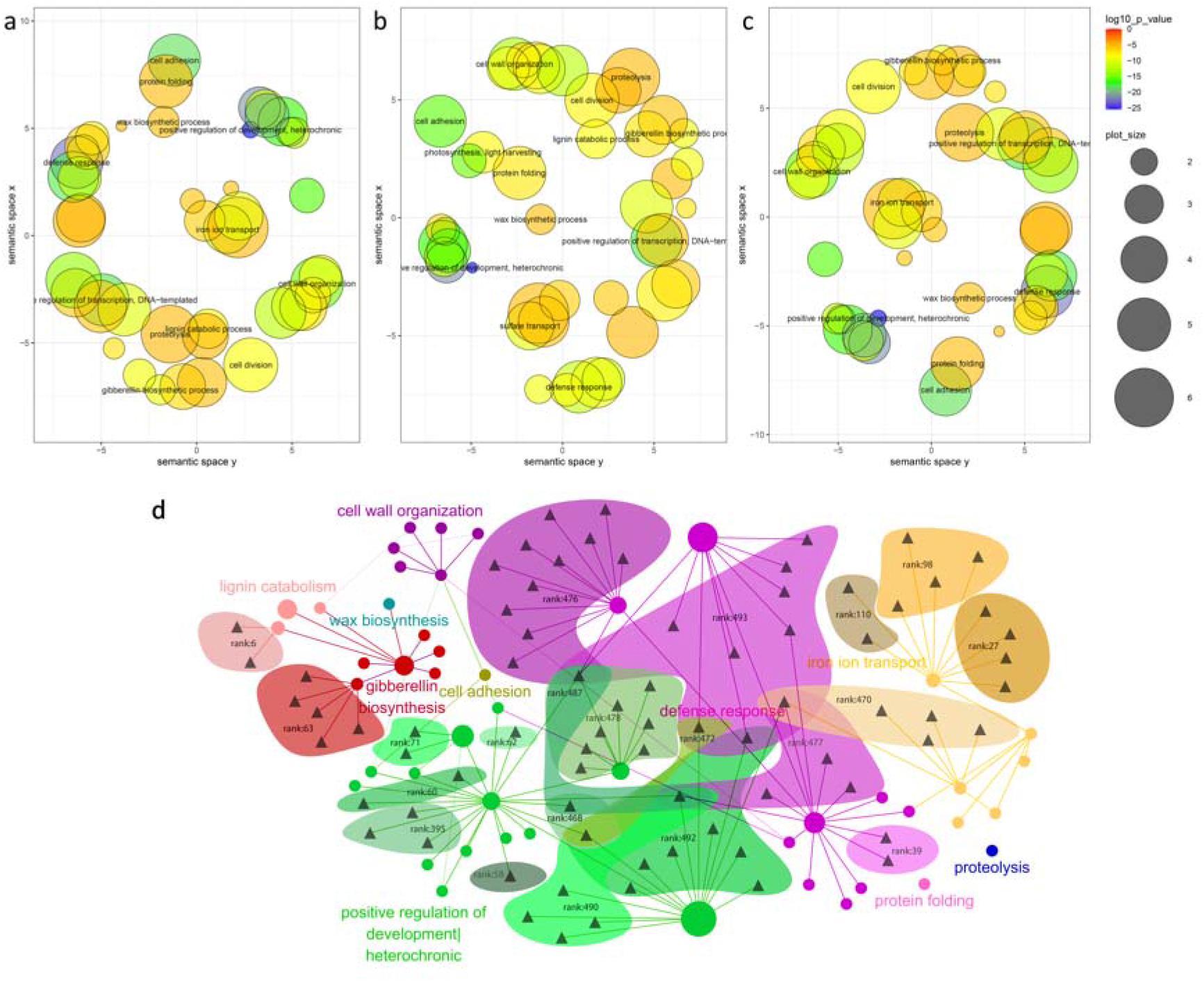
GO term analysis of MTRM genes under various abiotic stresses (a-c). (a) GO semantic correlation analysis of abiotic stress, (b) drought stress, and (c) salt stress, and (d) the GO BP regulatory network of cooperative miRNAs under abiotic stresses. Triangles represent different miRNAs and circles represent different GOs. The size of the circle is determined by the number of genes contained in the GO in this article. The color of the circle depends on the representative GO. The areas with different colors show the modules obtained by our method.

Subsequently, we constructed the GO BP regulatory network of cooperative miRNAs under soybean abiotic stress for the above main regulatory GO categories and miRNAs, as shown in Figure 5(d). Gene expressions related to abiotic stress responses are mainly regulated by multi-component miRNA families. For example, the miR167 family regulates the resistance response pathway; the miR171 family regulates the gibberellin biosynthesis pathway, while the miR395 family participates in regulating the iron uptake. Moreover, some miRNAs have multiple GO functional partitions, such as miR156b, which regulates developmental growth, the timing of developmental events, the response to hormones, and the response to heavy metal cadmium. The miRNA families and regulatory pathways involved in MTRM are detailed in Supplemental Dataset S4.

## Discussions

miRNAs are major regulators of plant growth and development. They can also regulate environmental responses [58-67]. Hence, the study of the role of miRNAs is crucial—not only to understanding the basic events of plant biology but to improve breeding for higher yields and more resilient crop plants. While a variety of papers have noted the role of one or a few miRNAs in regulating plant stress responses, a global analysis of the cooperative interactions is lacking. To study miRNA regulation in response to abiotic response in the soybean, we collected a large number of soybean MTIs. In addition, we collected MTIs from the model plant *Arabidopsis thaliana* and the legume *Medicago truncatula* and used them to predict potential targeting relationships in soybeans through homologous gene analysis. Meanwhile, psRNAtarget, TAPIR, and Targetfinder were used to support the targeting relationships. To ensure the reliability of the data, we mainly chose drought stress, salt stress, cold stress, and heat stress.

In the process of bi-clustering soybean MTIs, CUBiBit obtained many MTRMs based on targeting relationships between miRNAs and their target genes. To merge related MTRMs, we proposed a multi-level iterative fusion method of soybean MTRMs based on soybean gene networks. The method determines whether the miRNAs of the two modules have intersections, and the distribution of their genes in the soybean gene network determines whether they can be fused into a unified class.

We mined 483 soybean MTRMs, which provide a data reference for the analysis of the cooperative miRNA mechanism of the soybean. Some MTRMs are involved in the biosynthesis of chalcone, which is derived from the general phenylpropanoid pathway that plays a wide variety of roles in soybeans, as well as other plants. In most cases, gene regulation in each MTRMs involved a multi-component miRNA gene family. In some cases, these families were predicted to act cooperatively, which is consistent with the conclusion of Wang et al. [27]. The functions of a specific miRNA family can represent the main functions of miRNAs in the family, such as miR396 and its growth regulatory factors (GRFs) [68]; moreover, these regulatory factors include drought stress, high salt stress, low-temperature stress, and ultraviolet radiation stress. Interestingly, we found that miRNAs from different families are also involved in the same regulatory gene clusters, which indicates that different miRNA families may have cross-family cooperative regulatory mechanisms in regulating certain functions. In contrast, miRNAs in the same family can be in different MTRMs; for example, the miR171 family (miR172b-5p, miR172h-5p, miR172f, miR172g, miR172j, and miR172k) are in multiple regulatory modules during drought and salt stress. Such hub miRNAs may be useful research targets for exploring soybean resistance mechanisms and resistance to breeding research under different stresses. After further combining the analysis of differentially expressed genes in soybeans under various stresses, we obtained the miRNA-GO regulatory network under abiotic stress. The GO BP contains a variety of important related pathways for understanding the common mechanisms in stress response. The research covering the plant miRNA regulation module can analyze the coordination mechanism of miRNA from a global perspective and determine the regulation relationship between modules, which may help explore the regulation mechanism of soybean miRNAs.

## Methods

We collected soybean MTIs from *Arabidopsis thaliana* and *Medicago truncatula* databases and publications on miRNAs and genes of soybean response to several abiotic stresses. Subsequently, we used homology prediction on the collected MTIs to expand the soybean MTIs. We used the biclustering method to mine the soybean MTRMs to perform overlap analysis on the results and practiced redundancy removal. Then, based on the soybean gene interaction network, biclustering was applied through multi-level iteration. Finally, based on soybean abiotic stress-related miRNAs and genes, the fusion regulatory module was screened to obtain soybean abiotic stress-related MTRMs. Figure 6 shows a flowchart of our tasks and results.

**Figure 6.**
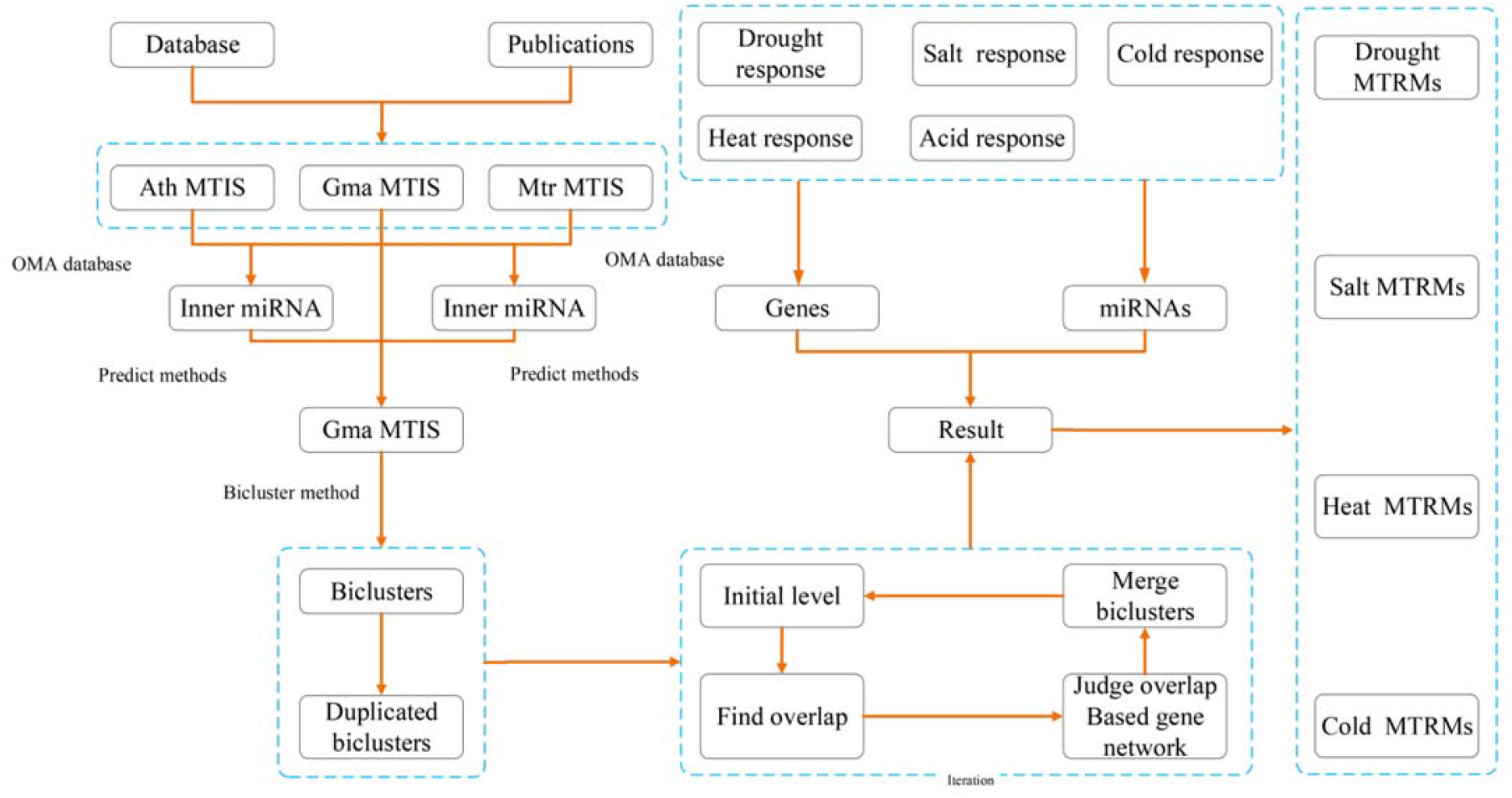
Flowchart of the authors’ research method.

### Data collection

We collected miRNA-target data of *Arabidopsis thaliana*, soybean and *Medicago truncatula* based on experimentally verified degradome sequencing results from databases (DPMIND, (c) Tarbase, mirTarbase, and Starbase [93-96]) and publications (Supplemental Table 1). In addition, we collected the miRNA information of the three species from the miRbase [97], the gene annotation of the species in the NCBI, EnsemblPlants, and the Phytozome [98, 99]. We also downloaded the homologous genes of *Arabidopsis thaliana* and *Medicago truncatula* in Orthologous MAtrix (OMA) [100]. Besides, we downloaded the soybean cDNA sequence and soybean gene GO annotations from SoyBase [101], and obtained soybean gene network data from SoyNet [102].

### Data processing

Due to the differences in the annotation methods in various databases and the differences in the data formats in the publications, we unified the miRNA and gene formats in the data and put the data of the same species together. We annotated the miRNA-target data based on the collected and processed miRNA details and the gene annotations derived from the data of three species, including miRNA target data, related notes and data sources. After processing the duplicated data, we obtained the miRNA-target data of the three species.

### Homologous extension

We chose *Arabidopsis thaliana* and *Medicago truncatula* to explore the potential targets. *Arabidopsis thaliana* as a model plant has rich high-quality data. *Medicago truncatula* and soybean are closely related and have many similar biological characteristics. We extracted the miRNA sequence and removed redundant miRNAs with the same sequence in the soybean and *Arabidopsis thaliana*. Subsequently, we extracted the target gene corresponding to the miRNA ID. Based on these targeted genes, we obtained soybean genes homologous to these genes from the *Arabidopsis thaliana*-soybean homologous genes downloaded by OMA. We assumed that if the sequences coexist and the genes are homologous, targeting relationships may exist. Therefore, these homologous genes may be targeted by these miRNAs in soybeans.

Targets obtained only based on homology information may not exist; so, we extracted these miRNA sequences and the cDNA sequence of target genes (SoyBase) and used miRNA-target prediction tools to predict potential relationships. Most of the existing plant miRNA target gene prediction tools showed high accuracy in *Arabidopsis thaliana*, but the prediction results in non-*Arabidopsis thaliana* plant data were different. Therefore, we chose psRNAtarget, TAPIR, and Targetfinder, whose results were better in non-Arabidopsis plants to predict potential soybean miRNA-target relationships [104]. The three prediction software tools have different scoring methods. We analyzed their respective scores and merged them. The homology extension method for *Medicago truncatula*-soybean is the same as above.

### Clustering method

The current research on miRNA targeting relationships is mostly based on one-to-one relative targeting. However, the miRNA targeting relationship is a complex interaction. The traditional clustering method is to cluster the same type of data, such as k-means, whose mining results in the miRNA-target regulatory module are poor because the targeting of miRNAs is sparse. The relationship between miRNA and the target gene is a bipartite graph structure; thus, the miRNA-target regulatory group can be found by analyzing the bipartite graph. Bimax has been widely used in gene expression data [103]. However, for more data, the calculation cost is higher. Domingo et al. proposed the BiBit [69], whose results are almost the same as that of Bimax, but the calculation time is still longer. Subsequently, ParBiBit and CUBiBit were proposed [57], which shortened the computing time and provided an optimized method for finding modules in larger data. We added the miRNA-target data based on the homology expansion predictions obtained from *Arabidopsis thaliana* and *Medicago truncatula* into the collected soybean miRNA-target data and extracted the miRNA-Target data with GO annotations and glyma2ID based on the soybean gene annotations of SoyBase. We used the CUBiBit to perform bi-clustering to obtain the results.

### Overlap and iterative fusion

The primary result obtained by CUBiBit was mostly a fully-connected bipartite graph. However, the relationship between miRNA and target gene is complex and interactive. Therefore, we proposed a method of iterative fusion for MTRM modules based on a gene interaction network (Figure 7).

**Figure 7.**
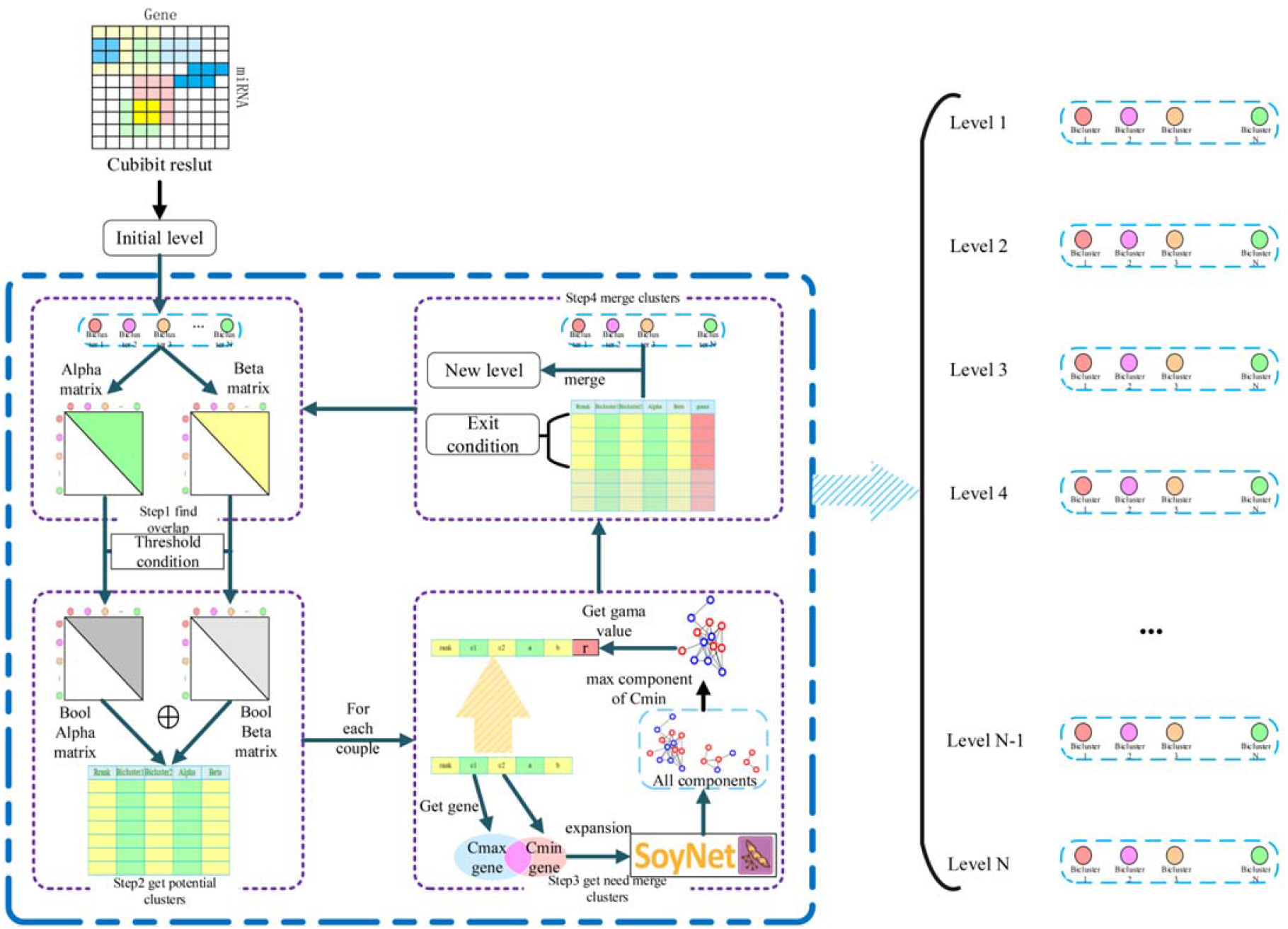
MTRM iterative merge algorithm flowchart used to derive a gene interaction network.

We detected completely included classes in the clustering results and removed the included classes as the initial level result. First, for each class of this level containing miRNAs and genes, we judged the degree of overlap with other classes of miRNA and genes to form alpha and beta matrices, both of which are upper triangular matrices. After that, we set two thresholds of miRNA and genes that can be potentially merged for the two classes. We then recorded the two classes that met the potential fusion class-class table requirements to form a Boolean matrix. The initial threshold of alpha was 0.3, and each iteration increased at a pace of 0.05 to conservatively determine the fusionable module and to keep this value unchanged after increasing to 0.8. It was sufficient if the beta threshold was greater than 0. Next, we extracted the union of two class genes, and the network blocks of this pair of genes with a depth of 2 layers based on the SoyNet network, for each pair of classes in the potentially merged class-class table. Subsequently, from the obtained block set, network blocks containing smaller classes were extracted. We assumed that the network block with the most smaller class genes represented the function of the genes in the smaller class. Therefore, judging the number of genes in the major category of this network block can determine whether the genes of the two categories are similar in function. If the genes of the two classes were concentrated on a network block, which means that their genes interact closely and meet the conditions of potential fusion, the two classes can be merged. We compared the number of genes in the major category with the number of genes in all major categories in the sub-category function module to obtain scores and determine the correlation. Finally, we compared the number of genes in the major category with the number of genes in all major categories in the sub-category function module to obtain scores to determine the correlation. The threshold was recorded as gamma. When gamma > 0.3 was satisfied, the two classes are merged; otherwise, they would not be merged. For the class pairs that meet the fusion condition, we arranged them in descending order of alpha value and performed top-down non-repetitive fusion. Each class can only merge at most one class in one iteration. A new class set was formed as the new level, and the fusion result was the output. The next iteration would be performed and then another iteration until no fusion class pair could meet the two conditions.

### Result analysis

For the results of the above iterative fusion, the enrichment of the classes in each level were separately analyzed. For a bicluster, we extracted its genes, used SoyBase’s GO BP and GO MF for enrichment analysis, and used the corrected GO ID with the smallest p-value as the best enrichment result for this type of cluster. When evaluating each class, only the smallest p-value was not enough to judge the importance of the class. Here, we should observe the situation in all the GO IDs enriched by this class to make a reasonable judgment on this class. Therefore, we used the cluster score to evaluate the enrichment of the class. For all the enriched GO IDs of this class, we screened all the results with a p-value of less than 0.05 and then used Eq. (1) to calculate the cluster score of the class.

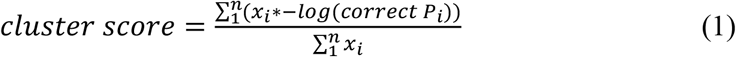

Among them, *n* is the number of gene ontologies enriched in the module, *x*_*i*_ is the number of genes enriched in the *i*-th GO, and *correct P*_*i*_ is the adjusted p-value of the *i*-th enriched GO.

### Abiotic Stress response MTRM

We collected the miRNAs of soybeans that respond to drought, salt, acid, and low temperature based on our studies of publications [22, 48, 105-121]. At the same time, we collected the differentially expressed soybean genes under various stresses and screened these genes with foldchange ≥ 2 and t-test p-value less than 0.05 as related genes under abiotic stress. Then we marked the genes in the module and calculated the p-value related to abiotic stress based on the hypergeometric distribution. Finally, we screened based on the cluster score calculated by the module, the p-value related to stress, and the proportion of miRNA related to stress. In addition, the screening procedures related to drought and salt stress were consistent with the screening steps of the abiotic stress module.

### Construction of miRNA-GO network under abiotic stress

Based on the results of MTRM mining under stress, we first screened the GO of the enrichment results in the screened module by p-value to remove the GO with a p-value less than 10^−5^; then, we performed a REVIGO semantic relevance analysis and extraction of concentrated representative GO channels. Based on the MTI data, the miRNA-GO relationship data was constructed through the gene pointed to by the miRNA in the module, and the enriched go pathway to which the gene belongs. The relationship between GO is based on the results of REVIGO and the GO similarity calculation. The relationship is presented by setting a threshold to remove some weaker relationships. More detailed parameters are provided here or in the location of the specific figure.

## Supplemental data

**Supplemental Table S1**. Summary of ATH, GMA and MTR MTIs sources.

**Supplemental Table S2**. Potential targets found by homology expansion.

**Supplemental Table S3**. Comparison of the biclustering results of soybean MTRMs.

**Supplemental Table S4**. The top 5 GO enrichment of each top 5 soybean MTRMs.

**Supplemental Table S5**. Top 5 soybean abiotic response MTRMs.

**Supplemental Dataset S1**. miRNA function annotation.

**Supplemental Dataset S2**. Screening results of soybean MTRMs related to abiotic stress, drought and salt stress.

**Supplemental Dataset S3**. Stress-specific MTRMs and shared MTRMs.

**Supplemental Dataset S4**. GO terms of miRNAs in the soybean abiotic response MTRMs.

**Supplemental Figure S1**. Potential soybean MTIs predicted based on three prediction tools. (a) 961 pairs of MTIs obtained based on the Arabidopsis data, (b) 986 pairs of MTIs obtained based on the Medicago data, and (c) 1,189 soybean MTIs from the union between (a) and (b).

**Supplemental Figure S2**. The differentially expressed genes in each MTRM under other stress after the screening.

## Acknowledgments

We would like to thank Ms. Carla Roberts for thoroughly proofreading this paper.

